# Optimal sparse olfactory representations persist in a plastic network

**DOI:** 10.1101/528125

**Authors:** Collins Assisi, Mark Stopfer, Maxim Bazhenov

## Abstract

The neural representation of a stimulus is repeatedly transformed as it moves from the sensory periphery to deeper layers of the nervous system. Sparsening transformations are thought to increase the separation between similar representations, encode stimuli with great specificity, maximize storage capacity as associative memories, and provide an energy efficient instantiation of information in neural circuits. In the insect olfactory system, odors are initially represented in the periphery as a combinatorial code with simple temporal dynamics. Subsequently, in the antennal lobe this representation is transformed into a dense spatiotemporal activity pattern. Next, in the mushroom body Kenyon cells (KCs), the representation is dramatically sparsened. Then in mushroom body output neurons (MBONs), the representation takes on a new dense spatiotemporal format. Here, we develop a computational model to simulate this chain of olfactory processing from the receptor neurons to MBONs. We demonstrate that representations of similar odorants are maximally separated, measured by the distance between the corresponding MBON activity vectors, when KC responses are sparse and that the sparseness is maintained across variations in odor concentration by adjusting the feedback inhibition KCs receive. Different odor concentrations require different strength and timing of feedback inhibition for optimal processing. Further, *in vivo*, the KC–MBON synapse is highly plastic, and changes in synaptic strength after learning can change the balance of excitation and inhibition and may lead to a change in the distance between MBON activity vectors of two odorants for the same level of KC population sparseness. Thus, what is an optimal degree of sparseness before odor learning, could be rendered sub–optimal post learning. Here, we show, however, that synaptic weight changes caused by spike timing dependent plasticity increase the distance between the odor representations from the perspective of MBONs and do not lead to a concomitant change in the optimal sparseness.

**Author Summary:** Kenyon cells (KCs) of the mushroom body represent odors as a sparse code. When viewed from the perspective of follower neurons, mushroom body output neurons (MBONs), an optimal level of KC sparseness maximally separates the representations of odors. However, the KC–MBON synapse is highly plastic and may be potentiated or depressed by odor–driven experience that could, in turn, perturb the optimality formed by pre–synaptic circuits. Contrary to this expectation, we show that synaptic plasticity based on spike timing of pre- and postsynaptic neurons improves the ability of the system to distinguish between the representations of similar odors while preserving the optimality determined by pre–synaptic circuits.

## Introduction

The neural representation of an odor is transformed repeatedly as it traverses different layers of the olfactory system (1). Some transformations separate the representations of odorants to enable easy discrimination (2)(3). Other transformations prepare an odor representation for eliciting behaviors by associating it with other sensory inputs and providing the context necessary for action and memory (4). In the American desert locust, *Schistocerca americana*, neural networks upstream of the KC–MBON synapse appear to work best as pattern decorrelators while downstream circuits appear to be specially structured to encode associative memories and organize behaviors elicited by stimuli. The olfactory network, from the receptor neurons through the antennal lobe and on to the mushroom body, is largely feedforward, and odor representations are progressively decorrelated and optimized in several ways as they traverse these layers. Odor representations arrive at the MBONs via synapses that are highly plastic and may change with the dynamic olfactory milieu of the animal (5). Here, we ask, how does the olfactory network preserve an optimal odor representation despite activity–driven changes in the synaptic weights of the networks?

In the locust, olfactory processing in the nervous system begins when odorant molecules bind to receptors on neurons in the antennae. This leads to the opening of ion channels and a cascade of events that can lead to spiking, the suppression of spontaneous firing, or simple sequences of excitation and inhibition. Olfactory receptor neurons can be tuned narrowly or broadly, firing vigorously for some odors and less so or not at all for others (6,7); thus, the identity of responsive neurons helps encode the stimulus. Temporal features of receptor neuron spiking, including response latency and duration also contribute to encoding the identity of the odor (8). Olfactory receptor neurons provide input to excitatory PNs and local inhibitory (and likely some excitatory) interneurons in the antennal lobe. This dense network, with recurrent connections between excitatory and inhibitory neurons, transforms the odor representation arising in receptor neurons into a more elaborate spatiotemporal pattern (1,9,10) where the identity, concentration, and timing of the odor are represented by the identity of responsive PNs, the temporal structure of their spiking, and correlations across the PN population. Most PNs respond in some way to most odors (1), collectively providing a dense spatiotemporal representation of an odor. KCs in the mushroom body receive inputs from PNs and transform this dense representation into a sparse code (11) in which rare spikes occur with millisecond precision and great specificity, together describing the attributes of the eliciting odor. The sparseness of KC spiking is orchestrated by a combination of membrane conductances that ensure a high spike threshold, and feedback inhibition from a giant GABAergic neuron (GGN) proportional to the drive it receives from the full population of KCs (12). Thus, GGN adaptively regulates the output of KCs, maintaining its sparseness over a large range of odor concentrations. The successive transformations that odor representations undergo, from dense spatiotemporal to sparse activity patterns, are thought to progressively decorrelate and distinguish odor representations and prepare them for valence and motor decisions, and storage as memories (13). Convergent KC activity is read out by a relatively small number of MBONs. The KC–MBON synapse undergoes experience dependent plasticity (5) (see (14) for a similar circuit in Drosophila) in a form that can be modified by associating an olfactory stimulus with a reward (5,14) mediated by octapamine. Together these features mark the KC–MBON pathway as one where sparse, decorrelated odor representations are combined with input from a reward pathway (15).

Using a model network that simulates olfactory processing in the locust from receptor neurons to MBONs, we show that the distance between the representations of different odorants, measured as the distance between MBON activity vectors, is maximized for a particular level of KC response sparseness. The degree of sparseness is determined by transformations of the odor representation in circuits before the KC–MBON synapse. However, what level of sparseness is optimal for odor discrimination by MBONs is determined by the weights of KC–MBON synapses. KC–MBON synaptic weight is, in turn, subject to the animal’s experiences, mediated by octopamine reward. This gives rise to a potential conundrum: the degree of sparseness determined by the circuits prior to the KC–MBON synapse could be rendered sub–optimal by modulations to the weight of that KC–MBON synapse by associative learning of particular odors. Here, we explore how the olfactory system guards against this loss of optimal sparseness. We show that the spike timing dependent plasticity operating on the strength of KC–MBON synapse not only retains the value of optimal sparseness despite learning-dependent changes in synaptic strength, but further improves the ability of the olfactory system to differentiate between odors.

## Results

In this study we sought to address two questions. First, from the perspective of MBONs, is there an optimal value of coding sparseness to maximally separate odor representations? Second, if an optimal sparseness exists, does plasticity at the KC– MBON synapse alter it, making post–learning odor representations sub–optimal? To address these questions, we constructed a computational model of the locust olfactory system consisting of the olfactory receptor neurons, the antennal lobe network of PNs and local inhibitory interneurons, the KCs of the mushroom body, and a layer of MBONs (Figure 1a). The model antennal lobe network generates many of the key responses previously documented in the locust (16,17). The output from the antennal lobe diverged widely to an array of 15,000 model KCs. This pattern of connectivity has been hypothesized to help decrease the overlap between odor representations (18,19). KC output then converged onto a small group of MBONs. To establish whether an optimal value of sparseness exists, we systematically varied the sparseness of KC responses and checked the ability of MBONs to differentiate between two similar odorants. We then introduced spike timing dependent plasticity (5) in the KC–MBON synapse and simulated the network using multiple instances of randomly interleaved odorants to map the effect of synaptic plasticity on the optimal sparseness of KC responses.

**Figure 1.**
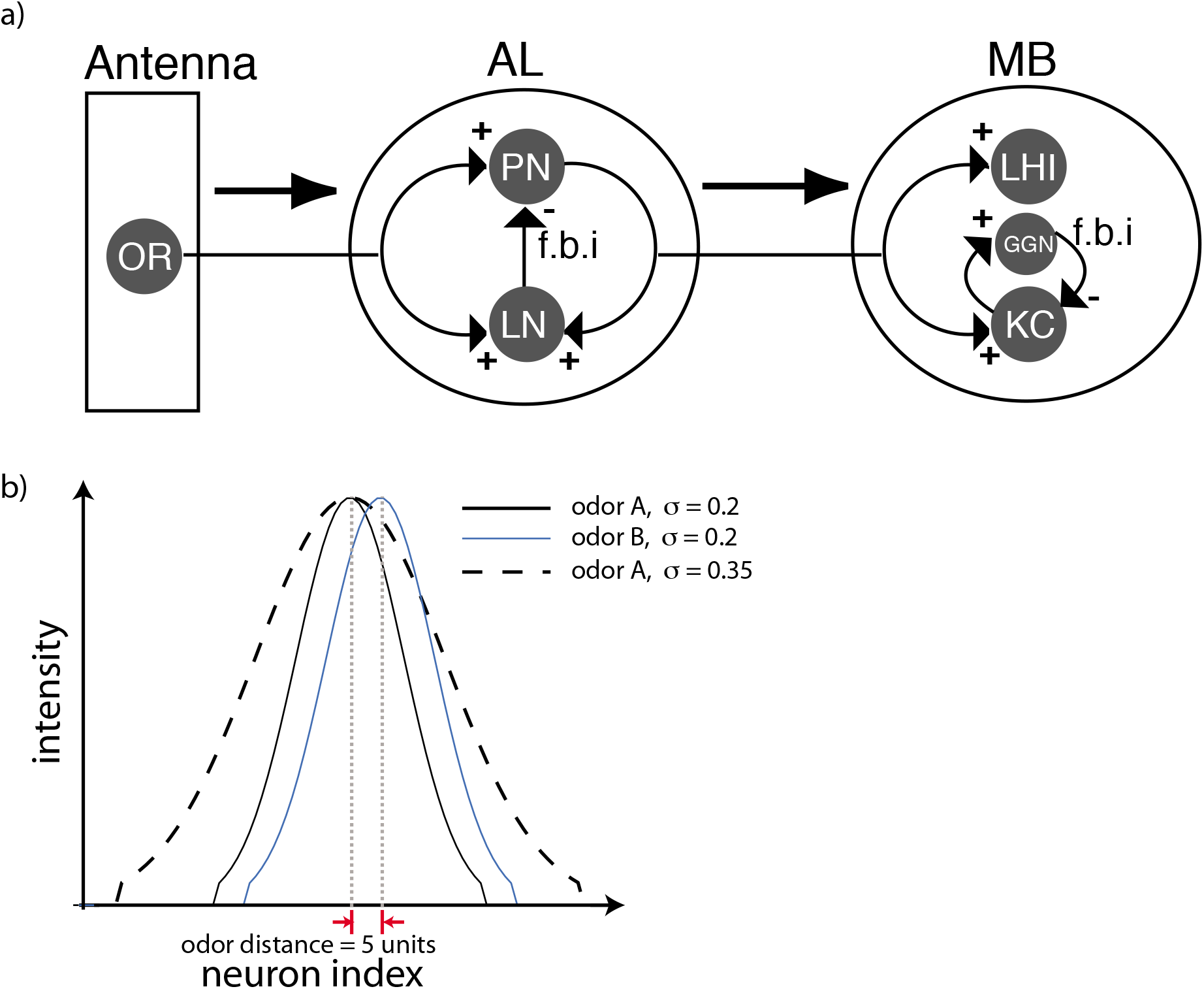
a) Schematic of the locust olfactory system. The first layer consists of olfactory receptor neurons (OR) in the antenna that provide input to PNs and local inhibitory interneurons (LNs) in the antennal lobe (AL). PNs project to KCs of the mushroom body (MB). In addition to excitatory input from the PNs the KCs also receive inhibitory input from the Giant GABAergic neuron (feed–back inhibition). b) Input to the AL. Each PN (LN) indexed on the x–axis receives a dc input of intensity shown along the y–axis. The concentration of the odor is characterized by the width of the function describing the input intensities while the identity of the odor is characterized by the position of its the peak.

### Coding sparseness determines the distance between odor representations

Odor input activates the ORNs that drive the neurons of the antennal lobe. We did not explicitly model the ORNs; rather, we simulated simplified ORN activity as depolarizing pulses to the PNs and interneurons (20–22). This input had an initial rise time of 60ms and a decay time of 200ms (Figure 2c bottom trace). The amplitude of the pulse was constant for 1000ms except for noise that was 5–10% of the amplitude of the pulse. Each odorant was defined by the subset of antennal lobe neurons it activated and the amplitude of the depolarizing input to each. In Figure 1b the amplitude of input to the PNs of the network is shown for two odorants (solid lines). The PNs were arranged such that the amplitude profile resembled a Gaussian curve. The input curve was set to zero when the amplitude decreased below a threshold value. Note that the arrangement of PN indices according to a Gaussian activation profile does not imply any spatial structure; that is, the neuron with index *i* need not be physically adjacent to neurons with index *i−1* or *i+1* since network connections were chosen randomly. However, by defining an odor in this manner, we could conveniently and continuously vary the identity and the concentration of odorants. The identity of an odorant could be varied by moving the location of the peak while the concentration could be increased by widening the Gaussian to recruit more PNs and interneurons (Figure 1b, dashed line) (23). We measured the responses of antennal lobe neurons to the odor input. As seen in earlier studies and in accordance with experiments done *in vivo*, the local field potential (measured in our model as the mean membrane potential of all the PNs) showed an odor-elicited 20Hz oscillation (Figure 2b). This global pattern was elicited by different odors and concentrations. As *in vivo*, during each cycle of the oscillation different groups of PNs were transiently activated. This spatiotemporal representation continuously changed over the course of the odor presentation due to mutual and transient inhibition between interneurons that, in turn, entrained the activity of PNs (20–22). Each odor–concentration pair that we tested generated a different spatiotemporal pattern. An example pattern of activity is shown in Figure 2a,c. The amplitude of the local field potential increased with increasing concentration (not shown here) indicating tighter synchrony between the projection neurons that spike during each cycle, consistent with earlier studies (23,24).

**Figure 2.**
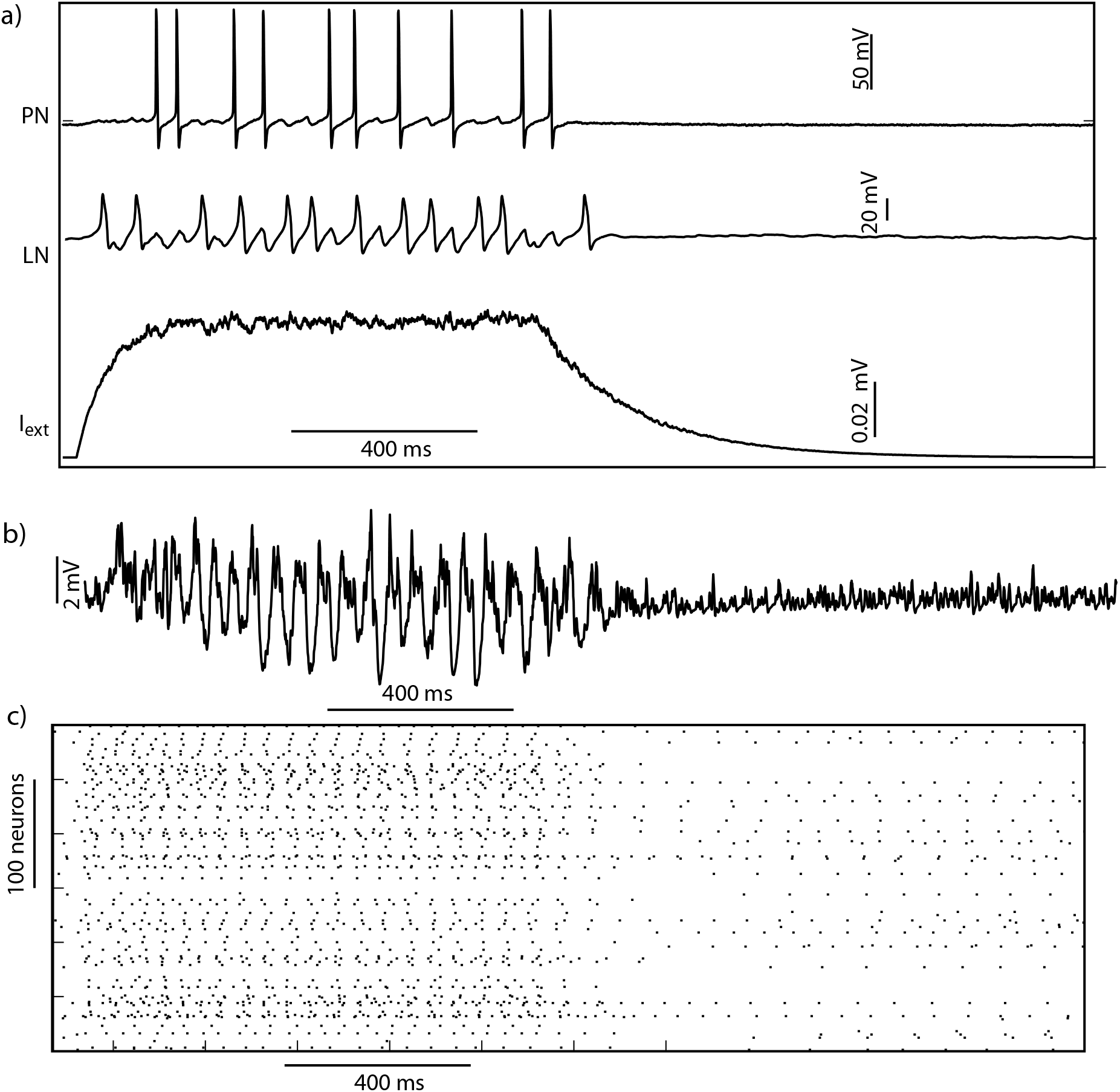
The top trace in (a) shows the response of a single PN to an external input (a, bottom trace) while the middle trace shows the response of an LN. The trace in (b) shows the summed projection neuron (PN) voltage response to an external input to the network. The input (I_ext_) (a, bottom trace) was scaled by intensity (shown in figure 1b) for each neuron. The PN raster plot for a reduced network with 300 PNs and 100 local neurons (LNs) is shown in c.

The PN responses were used as input to a group of 15,000 KCs. KCs are known to respond sparsely (few neurons fire rarely) to odor stimuli (11). However, the increased PN synchrony that accompanies increased odor concentrations (24) alone would lead to more densely spiking responses, disrupting the sparse code. How do KCs maintain sparseness over decadal variations in the concentration of the odor? Earlier studies hypothesized that input from PNs to KCs arrives along two pathways, a direct excitatory drive from the antennal lobe and slightly delayed feedforward inhibition from lateral horn interneurons (LHIs). Thus, cyclic pairs of excitatory and inhibitory input to KCs defined short windows of time during which KCs could integrate input from the antennal lobe (11). Notably, the duration of this window was dynamically modulated by changes in the concentration of the odor, which allowed KCs to fire sparsely despite large changes in the concentration (23). Recent work established that LHIs do not extend GABAergic projections to KCs, eliminating the possibility of feedforward inhibition (25). However, the cyclic inhibition underlying the dynamically modulated integration windows is generated by feedback from a single inhibitory neuron, termed the Giant GABAergic Neuron (GGN), which provides input to all KCs (12,16). Thus, cyclic inhibition regulates the sparseness of KC responses in an adaptive, concentration dependent manner (25,26). Given these earlier findings, we first sought to determine whether there exists an optimal value of lifetime sparseness (measured as the total number of spikes generated by all KCs during an odor presentation) to discriminate odors. To determine an optimal sparseness, if one such value existed, we needed to quantify the distance between odor representations from the perspective of downstream neurons that read KC output. KCs converge onto MBONs that generate distinct responses to odorants (27). Therefore, we used the MBONs as a read–out of KC responses. The Hamming distances between odor representations generated by MBONs were plotted as a function of different manipulations to the upstream network. To model the feedback inhibition from GGN that regulates the sparseness of KC responses using feedback inhibition, we drew on results from numerous studies showing that feedback inhibition in excitatory-inhibitory circuits mainly reduces later portions of the excitatory responses in each cycle. As inhibition strengthens, its onset occurs faster, thus reducing the excitatory response (23). This effect is self-limiting though, because excitation is needed to drive inhibition (26). Thus, rather than explicitly modeling GGN, we modeled the effect of feedback inhibition by selectively eliminating PN spikes that occurred after a thresholded phase of the LFP (Figure 3a). To do so we first filtered the LFP (40Hz) (red trace in Figure 3a) and calculated the instantaneous phase of the resulting 20 Hz oscillation using a Hilbert transform. Then we removed those spikes occurring beyond a threshold phase, denoted by *ϕ*, in each cycle of the LFP, shown by the shaded regions in Figure 3a. In this way we could directly control this threshold phase, and therefore artificially vary the window of integration and the sparseness of KC activity. As expected, as the window of integration widened, KCs received more input from PNs and generated progressively denser spiking activity. We plotted the density of KC spiking as a function of *ϕ* for different values of odor concentration (*σ*) (Figure 3c). Increasing odor concentrations led to denser KC responses for a given value of *ϕ*. To maintain a specific value of sparseness that maximally separated odor representations across a range of odor concentrations, the window of integration shifted to lower values as concentration increased through a mechanism that arises naturally from concentration–dependent changes in odor–response latencies in PNs (23,25)(Assisi et al., 2007; Gupta and Stopfer, 2012). To determine the separation between odor representations from the perspective of downstream targets, we used odor–elicited KC responses to drive a group of 100 MBONs (Figure 3b). Here the MBONs were modeled using a two–dimensional map that integrates pre–synaptic input and generates spikes in response to it (see methods). We binned the output of these neurons into temporal blocks, each block demarcated by the troughs of an LFP cycle. The response of MBONs during each cycle of an LFP oscillation was assigned one or zero to indicate whether it had spiked or not (Figure 3b).

**Figure 3.**
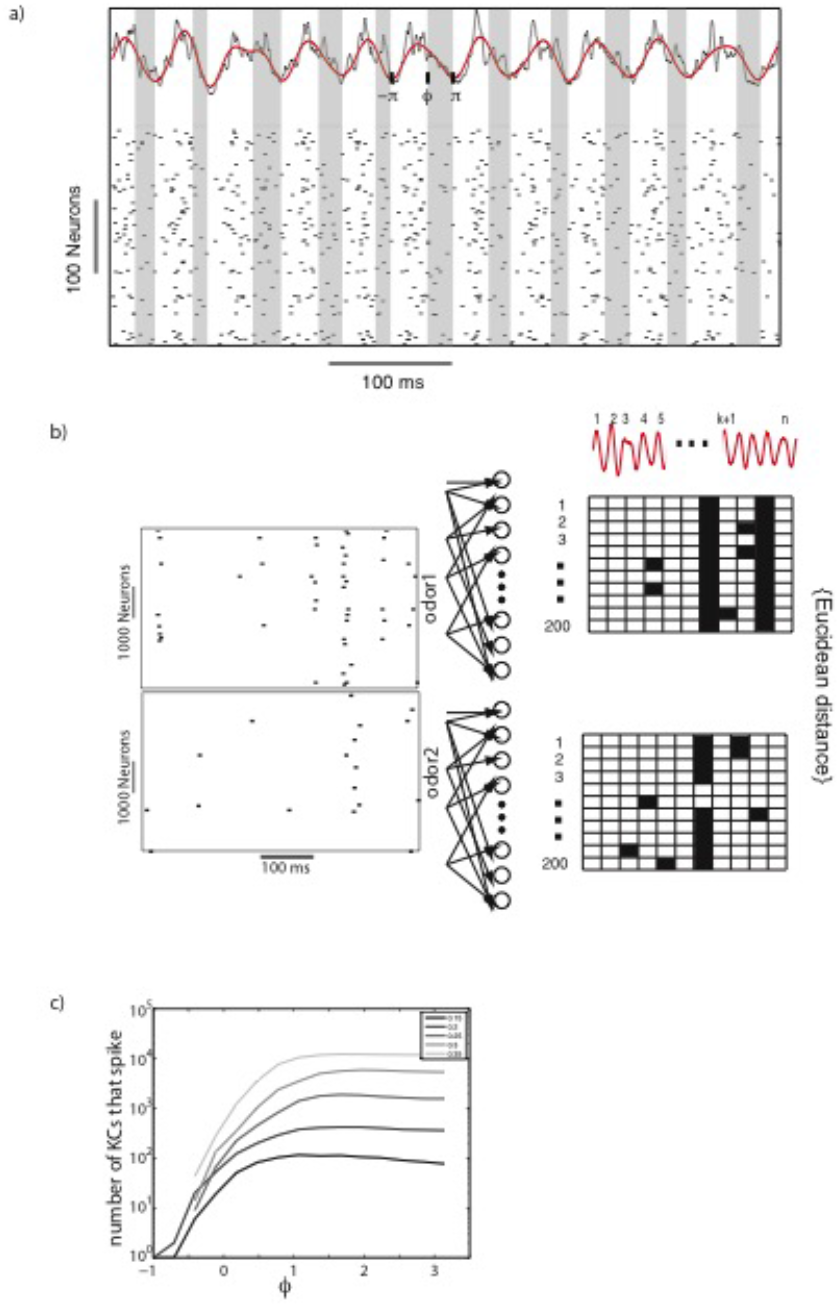
Individual KCs receive oscillatory input from the PNs in the AL. The sparseness of the KC responses was modulated by choosing a window of time over which each KC integrated input from PNs. That is, spikes occurring after a specified phase (*ϕ*), in the shaded region, were ignored and did not affect the KC responses. The raster plots in (b) show the responses of a subset of KCs to two different odors. These spikes were then fed to a layer of 100 beta lobe neurons. The response of the beta lobe neurons was converted into a binary spatiotemporal pattern. For each neuron, a single cycle of the PN-LFP was marked either 1 (dark) or 0 (blank box), depending on whether that neuron fired a spike during the cycle (Panels on the right in b). The Euclidean distance between these binary spatiotemporal patterns was used to calculate the distance between odors. The number of KCs that spike in response to an external input is plotted in (c).

We then calculated the Hamming distance between the representations of two odorants as a function of different windows of integration, as determined by *ϕ* (Figure 4). Since the density of KC spikes was a monotonically increasing function of *ϕ*, we used *ϕ* as a proxy for KC sparseness. We found that, for low odor concentration values (*σ*< 0.25), the peak distance between odor representations occurred when more PN spikes were allowed to affect KC responses throughout each LFP cycle (Figure 4a). Indeed, for low odor concentrations, despite the lengthy integration windows they evoked, the synchronization and thus density of input PN spiking remained low and the responses of KCs remained sparse. With increasing odor concentrations, we found that the peak shifted to lower values of *ϕ*. Furthermore, we observed a decrease in discrimination performance beyond a certain *ϕ* threshold. This occurred because KC responses became denser when the integration window expanded. Thus, for higher odor concentrations, we found a prominent single peak suggesting the existence of an optimal value of KC sparseness to maximize the distance between odor representations from the perspective of MBONs.

**Figure 4.**
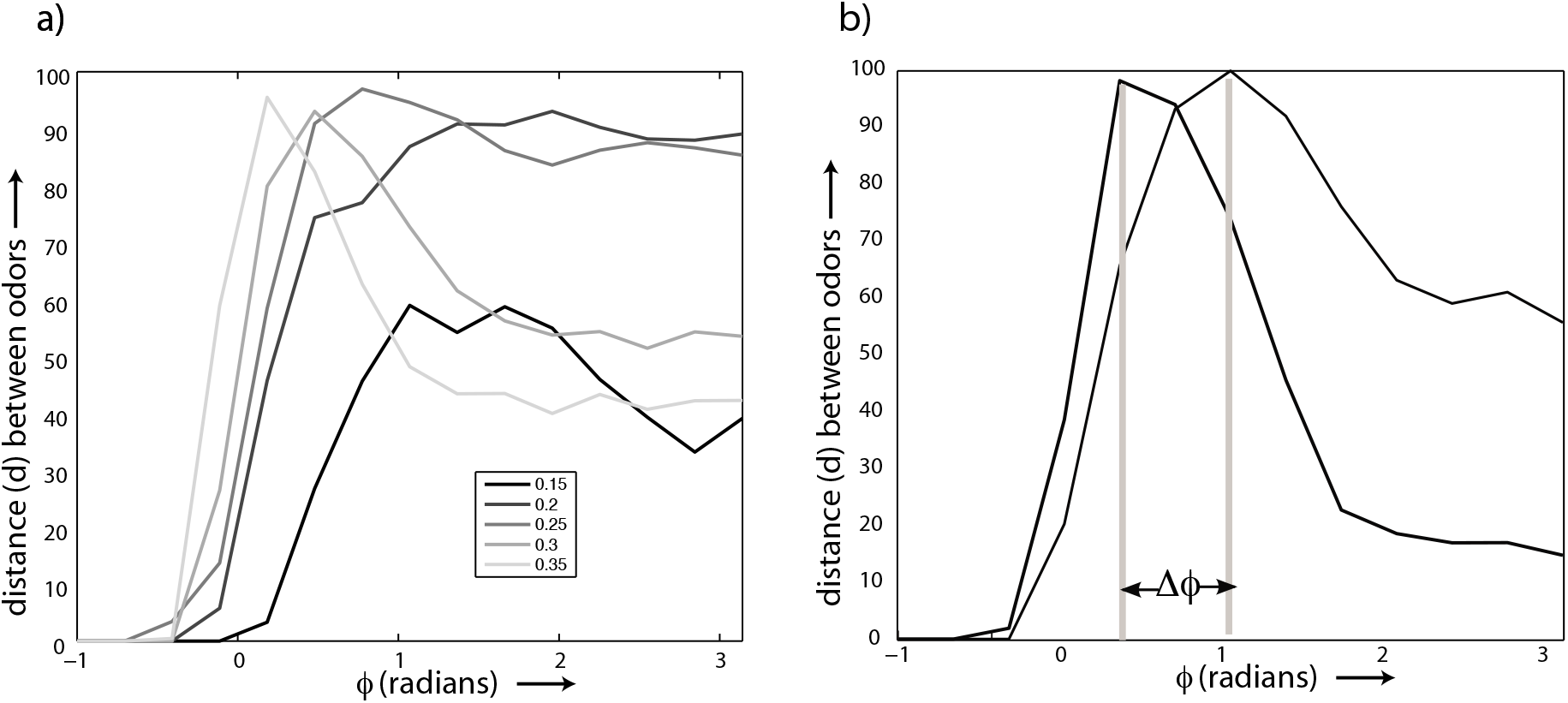
a) Distance between odors plotted as a function of the window of integration () for different values of the concentration (). b) Distance between odors for two different KC–MBON synaptic weights. The shift in phase of the peak distance is marked Δ*ϕ*.

Within a given animal the impact of KC spiking on MBONs can vary over time because the synapses linking them are plastic, changing in strength with experience (5). By amplifying or decreasing the impact of KC spiking, this synaptic plasticity has the potential to degrade the effective, optimized sparseness of the KC output, potentially affecting the distance between odor representations from the perspective of MBONs. To investigate this possibility, we systematically varied the weight of the input synapses to MBONs to determine how plasticity affects the distance between odor representations. We then simulated delivery of two similar odors of the same concentration by shifting the peaks of the distributions that characterized the two odors by 5 units with respect to each other, and for two different concentrations by adjusting the widths of the distributions (Figure 1b). As before, patterns of antennal lobe activity served as input to KCs that, in turn, drove MBONs. Here, we used the output of MBONs to measure the distance between odor representations for different values of KC sparseness. As before, here the MBONs were modeled as simple map–based neurons, summing the input they received from KCs and generating a spike in response to supra–threshold inputs. We then increased the weight of the KC–MBON synapse by a factor of 2. This manipulation led to a shift of the peak towards lower values of *ϕ* (Figure 4b). These simulations confirmed that the effective sparseness of KC output could change when the animal is exposed to different sets of odors that trigger plasticity in the KC–MBON pathway. This departure from optimal sparseness could be detrimental to subsequent circuits that, to appropriately inform behaviors, depend on an accurate distinction between odors.

### Spike timing dependent plasticity increases the distance between odor representations and preserves optimal KC sparseness

KC–MBON synapses appear to be powerful: a KC spike generates an EPSP in MBONs that is, on average, nearly an order of magnitude larger than EPSPs generated in KCs by PN spikes (5). Previous work established that the KC–MBON synapse undergoes spike timing dependent plasticity (STDP): potentiation when the presynaptic neuron fires before the post synaptic one, and depression when the presynaptic neuron fires after the post synaptic one (figure(6b) (5). This plasticity has been shown to maintain the oscillatory parcellation of information that begins at the antennal lobe and cascades all the way down to the MBONs. How does this plasticity affect the distance between odor representations when viewed from the perspective of MBONs?

To address this question, we modeled STDP in the KC–MBON synapse using a simple phenomenological model (Fig. 5b) (28,29). Following STDP rules, the model effectively modified the weight of the synapse depending on the time of occurrence of the presynaptic and the postsynaptic spikes such that each occurrence of a presynaptic spike before the postsynaptic spike led to an increase in the synaptic weight, and a presynaptic spike after the postsynaptic spike led to a decrease in the synaptic weight (see Methods for implementation details). We modeled MBONs parsimoniously as reduced spiking neuron models represented by two dimensional maps (30,31) (see Methods for details of implementation). The increase/decrease in weights with each pre–post pair of spikes is shown in Figure 5a. We specified a minimum and a maximum value for the synaptic weights so that the response of MBONs extended over a wide range of sparseness values. The change in synaptic weights (*dw*) depended on how close the current weight of the synapse was to the maximum allowed synaptic weight. The STDP equations were modeled so that for large synaptic weights (*w_max_*) synaptic depression dominates over potentiation and vice versa for small synaptic weights (*w_0_*) (32). Using this form of STDP, we then wired the 15,000 KCs to a layer of 100 MBONs. Each MBON received input from a randomly selected group consisting of 60% of the KCs. For simplicity, we did not implement lateral inhibitory connections between MBONs that are thought to enhance the contrast of input received from Kenyon cells (5).

**Figure 5.**
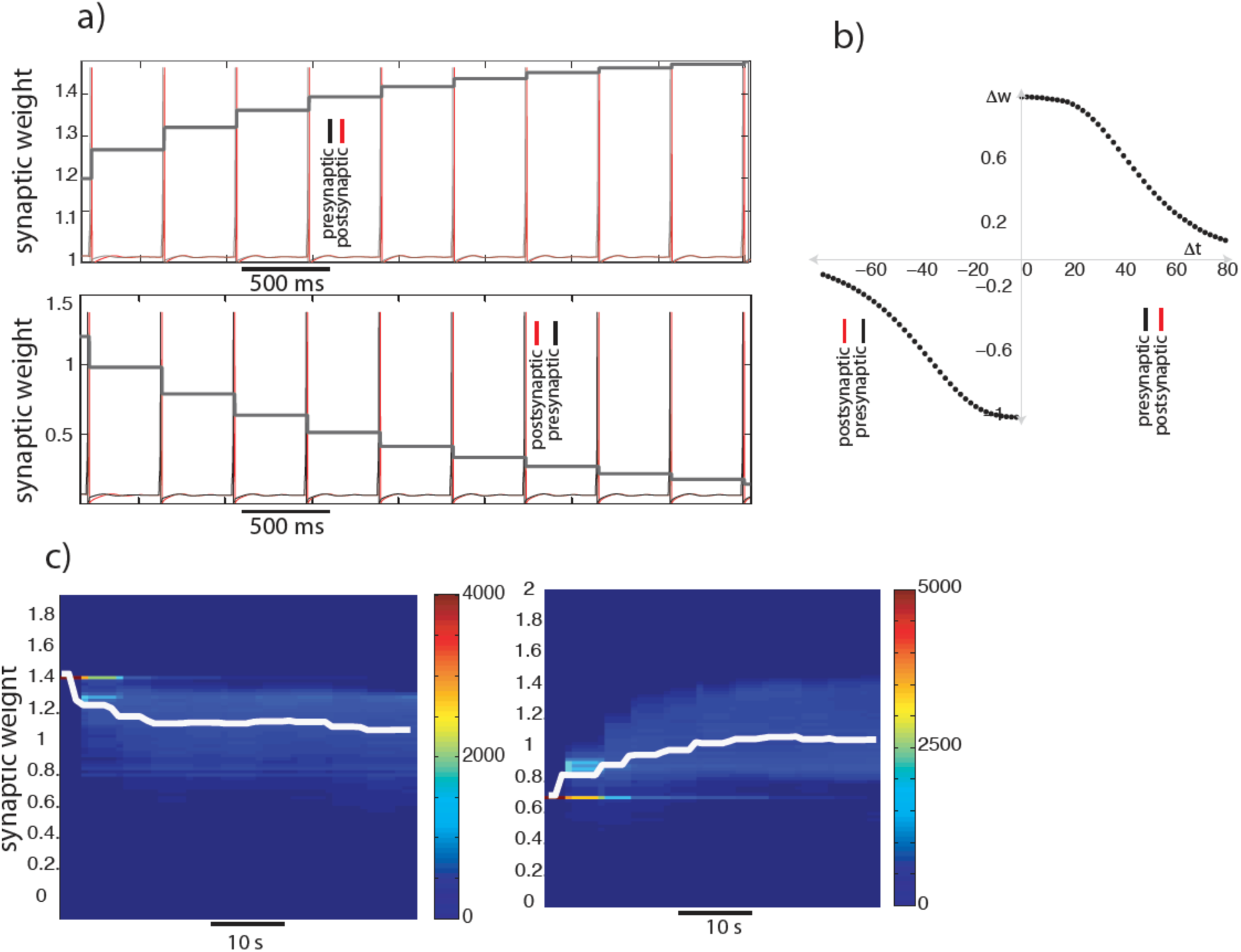
Spike timing dependent plasticity. The post–synaptic neuron (red) spikes follow that of the pre–synaptic neuron (black), leading to an increase in the synaptic weight (facilitation) (Fig. 5a top panel). The opposite temporal order (post–synaptic spikes occur before pre–synaptic spikes) leads to a decrease in synaptic weights (depression) (Fig. 5a bottom panel). The increase/decrease in synaptic weight (Δ*w*) is shown as a function of the time difference between the pre– and the post–synaptic spike (5b). When the presynaptic spike occurs before the post–synaptic spike Δ*t* is positive and otherwise negative. The distribution of synaptic weights of all KC-MBONs pairs evolves over time (c). In the left panel of (c) all the initial weights were set to 1.4. The system was then stimulated with different odors of varying concentrations. The weights were sampled at fixed intervals of time. The distribution of weights was plotted using a color map (see color bar for the frequency values). The mean synaptic weight was overlaid on the distribution (white trace). The right panel shows the temporal evolution of the synaptic weights when a low initial weight (0.6) was used.

To model odor stimulation, we randomly interleaved multiple instances of two similar odors (peak shifted by 5 units) and an odor that was different from these odors (peak shifted by 20 units) as input to the PNs. This simulated odor input evoked spatiotemporal patterns of activity in the antennal lobe that drove the KCs and the MBONs. Initially, the narrow distribution of synaptic weights of the KC–MBON synapses was, in separate simulations, centered around two different values (Fig. 5c, left vs right). Over successive odor presentations these synaptic weights changed. The median value of the synaptic weight is shown by the white lines in figure 5c. Since KC responses are very sparse, most of the weights did not change at all. Therefore, in subsequent analyses we chose a subset of weights that changed during the course of multiple odor presentations. We found that, over odor presentations, the distribution of synaptic weights evolved in a manner such that the median synaptic weight changed monotonically toward new value (approximately 1 in Fig. 5c). In our simulations we used two different initial weights distributions. Regardless of the specific initial weight distribution, we found that the median synaptic weight evolved towards the same value over multiple odor presentations.

We next investigated the evolution of the odor representation exhibited by MBONs concomitant with the evolution of the network weights. The peak distance between the representations of two similar odors increased as the weight distribution settled to its asymptotic values (Figures 67a and 67b; lighter colored curves correspond to the weights later in training). However, notably, the location of the peak marked as *ϕ_max_*, for which the distance between odor representations was maximum, depended on the STDP induced changes in the KC–MBON synaptic weight. KC sparseness (f) depended on the peak distance. Notably, the location of the peak did not change despite variations in the magnitude of the peak associated with weight changes caused by STDP. This effect may be seen in figure 7ca for two different concentrations of the simulated odorants. As odor stimuli were presented repeatedly to drive the network, the value of optimal sparseness remained the same as the network settled to its asymptotic weight distribution. This was true regardless of whether the initial weight distributions were set to values higher or lower than the asymptotic median value of the weight distribution post training of the network. At a low odor concentration (*σ* = 0.2) the changes in the distance curve (marked in progressively lighter shades) were small compared to the changes at higher concentrations (*σ* = 0.35, red curves in figure 7a). In all cases, however, the optimal degree of sparseness provided by the circuits before the KC–MBON synapse remained optimal after STDP mediated changes of KC–MBON projections. Thus, we conclude, an effect of STDP is to improve the ability of the olfactory system to differentiate between odors (Figure 6b) while retaining the value of optimal sparseness despite learning dependent changes in synaptic weight (Figure 6c).

**Figure 6.**
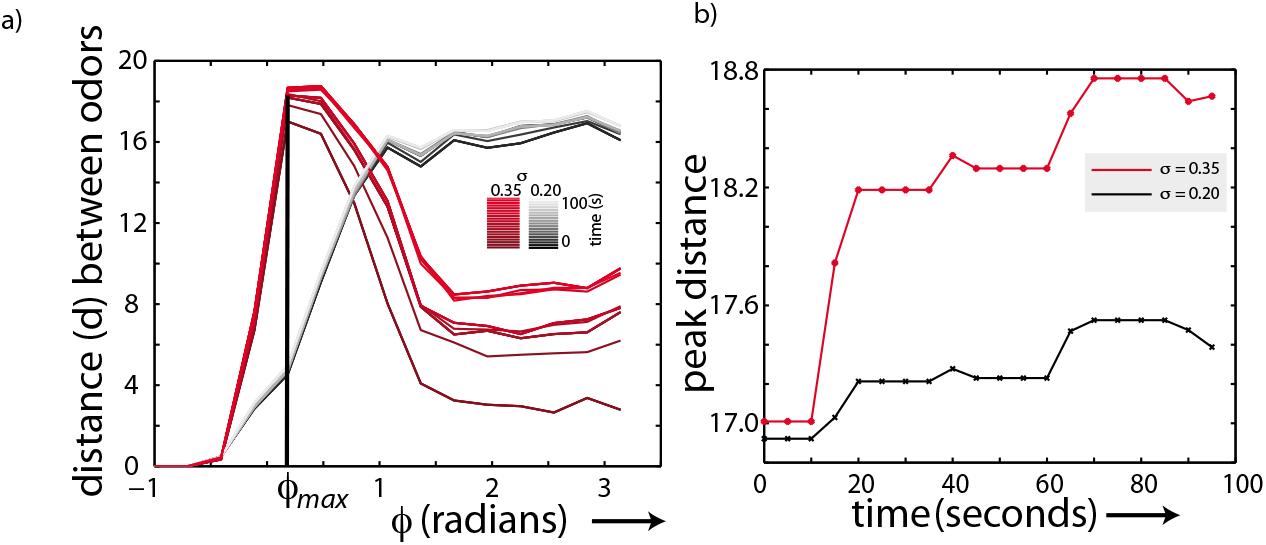
Role of STDP in odor discrimination. Distance between odors plotted as a function of the window of integration of KCs spikes (*ϕ*) for high (*σ* = 0.35 red) and low (*σ*= 0.2 gray) concentration values (a). Spike timing dependent plasticity reshapes the weights of the KC-beta lobe connections over time. Each line in the plot shows the odor distance–*ϕ* relationship at different snapshots in time ranging from 0 to 100 seconds. Darker shades indicate earlier times. (b) shows the maximum distance between two similar odors for high (*σ* = 0.35 red) and low (*σ* = 0.2 gray) concentration values over the time that STDP re–weighted the KC–MBON connections. Note, that the *ϕ* value maximizing the distance does not change during STDP-mediated learning.

## Discussion

In *Drosophila* and in locust, sparseness in KC firing is achieved by intrinsic high firing thresholds and feedback inhibition from a single neuron in each lobe, the anterior paired lateral (APL) neuron in *Drosophila* (33,34) and GGN in locust (12,16,25,27). This simple architecture, with a single neuron exerting outsized influence over the olfactory system, allows relatively simple experimental perturbations that selectively change the sparseness of KC responses. Indeed, as KC spiking increases in density, the ability of insects to differentiate between similar odors decreases, but the ability to differentiate dissimilar odors is not affected (34). This observation suggests that decreased sparseness increases the overlap between representations, but that KC representations of dissimilar odors are sufficiently distant and continue to be separable even when sparseness is compromised.

Here, we developed a model that couples multiple layers of the locust olfactory system. Our results demonstrate that olfactory circuitry drives odor-driven responses in KCs to a specific optimal value that, from the perspective of downstream neurons that receive convergent input from KCs, the MBONs, maximally separates odor representations. We hypothesized that changes in synaptic weights caused by experience-dependent plasticity could degrade what had been an optimal representation. However, our simulations show that, despite STDP-induced changes to the strength of the KC–MBON synapse, the value of optimal sparseness was maintained. This finding is particularly significant because the overall feedforward architecture of the insect olfactory system, featuring a near absence of feedback across layers, implies that downstream layers cannot ‘error-correct’ upstream representations. Thus, the connectivity between layers must assure that an optimal representation constructed in one layer continues to be optimal from the perspective of subsequent layers. Our model allowed us to explore the mechanism underlying the maintenance of optimal sparseness across circuit layers and despite neural plasticity.

In the locust olfactory system, odor representations are parceled into cyclic 50ms packets of information, a process that begins in the antennal lobe and cascades at least two synapses forward to the MBONs. This parcellation is maintained and stabilized against noise and other corruption by STDP that adjusts the strength of synapses when pre–synaptic input leads or lags post–synaptic output by tens of milliseconds (within an oscillatory cycle). Lateral inhibition across MBONs may further sharpen the odor representation. In this study, we employ a specific STDP rule where potentiation occurs when the pre–synaptic spike occurs before the post– synaptic response. When a reinforcing octopamine signal is delivered locally to the mushroom body a few seconds after a pre-post pairing has initiated STDP, the specific synapses that have participated in the pairing undergo an associative change (35) that consists of a transformation of the plasticity rule governing the dynamics of the synapse. Thus, STDP appears to play two roles. In the absence of reinforcement, it supports the oscillatory dynamics inherited from the antennal lobe by regulating the phase of MBON spikes. When reinforced with octopamine, the STDP rule for odor specific KC–MBON pairs may itself undergo a qualitative change STDP tags the synapses between odor specific KC–MBON pairs. Thus, the presence of a diffuse reinforcing signal can initiate highly specific changes. Our model pertains to the pre–reinforced KC–MBON synapse. This mechanism progressively increases the distance between odor representations while retaining the optimal sparseness at each processing level. We note that our analysis addresses a specific form of plasticity exhibited by the KC–MBON synapse, and may not generalize to other forms of plasticity that may be present.

The KC–MBON junction may be the location where the imperative of insect olfactory system changes from identifying the odor to associating the odor with other sensory and reward inputs. If so, MBONs may not require a precise representation of the odor. In fact, in *Drosophila*, the MBONs have been shown to be broadly tuned, and thus instantiate a representation more redundant than that of the population of narrowly tuned KCs. However, studies in locust have shown that the odor-elicited responses of MBONs, though densely spiking, are sensitive to the temporal ordering of KC input (27), and contain information about odor identity (36). In our analysis we included the dynamics of MBONs throughout the odor stimulus, parsing its spike trains into 50 millisecond bins (equivalent to one oscillatory cycle in locusts) and calculating the ability of the system to discriminate odorants over the entire duration of the odor. Our study suggests that odor representations are maximally separated when the neural representation of the odor in the mushroom body is optimally sparse. Despite challenges, the olfactory circuit of insects maintains this optimal sparseness over variations in the concentration and experience dependent plasticity.

## Methods

The model antennal lobe consisted of a scaled–down network of 350 PNs and 100 inhibitory interneurons (the locust antennal lobe contains roughly 800 PNs and 350 local neurons). Each neuron was modeled as a single compartment with voltage and calcium dependent currents with Hodgkin-Huxley kinetics. PNs generated *Na*^+^spikes while inhibitory interneurons generated *Ca*^2+^ spikelets, as seen in the locust olfactory system (36). The model of inhibitory interneurons included a *Ca*^2+^ current (*I_Ca_*) and a *Ca*^2+^ dependent *K* current that caused spike rate adaptation. Model PNs included a fast sodium current *I*_Ca_, a fast potassium current *I_K_* (37), a transient potassium A-current *I_A_* (38), and a potassium leak current *I_KL_*. The equations governing the dynamics of the neurons are as follows,

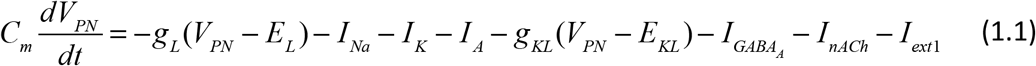

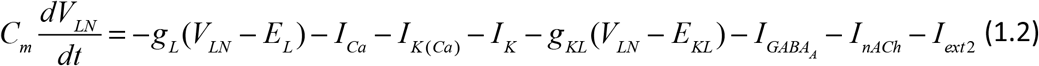

The passive parameters of the model were set as follows. *C_M_* = 1.43 × 10^−4^ *μS*, *g_L_ =* 0.15*μS* and *g_KL_* = 0.05*μS*. *E_L_* = −55*mV* and *E_KL_* = −95*mV*. The passive parameters were set the same for both the PNs (subscript *PN* in all the equations) and the inhibitory local interneurons (subscript *LN* in the equations). The intrinsic currents governing the dynamics of each neuron is given below.

Sodium current *I_Na_* is given by,

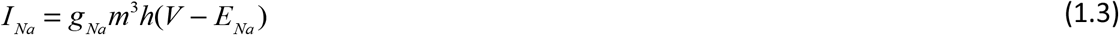

where, the *Na* conductance, *g_Na_* = 50*μS* and the reversal potential, *E_Na_* = 50*mV*. *m* and *h* are the activation and inactivation variables that are given by,

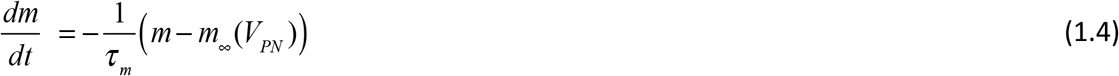

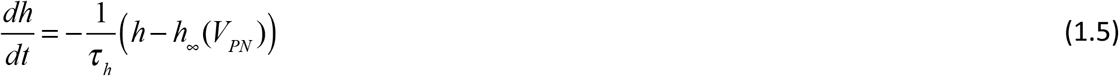

where, 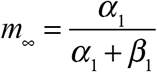 and 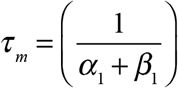 with 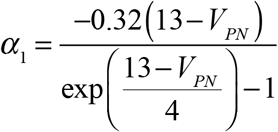 and 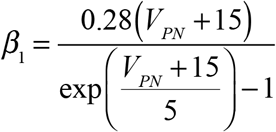. The steady state values of the inactivation variable *h* and the time constant *τ_h_* are given by 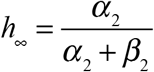 and 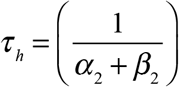, where, 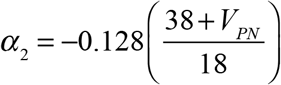 and 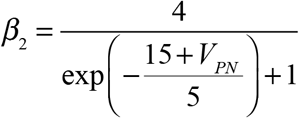

The equations describing the potassium current *I_K_* for both PNs and inhibitory interneurons are as follows,

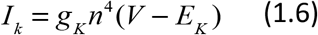

where, *g_K_* = 10 and *E_K_* = −95. The activation variable of the *K* current is given by,

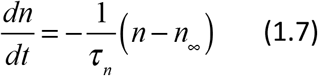

where, 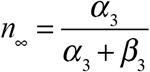 and 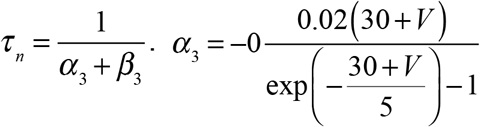 and 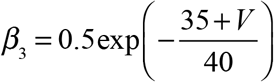

A transient potassium current, *I_A_*, in PNs was described by the following equation,

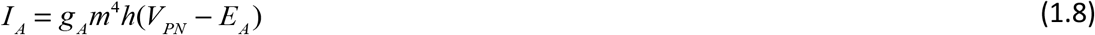

where, *g_A_* = 10*μS*. The steady state values of the activation and the inactivation variables are given by 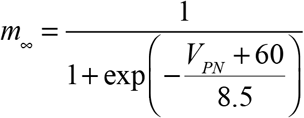 and 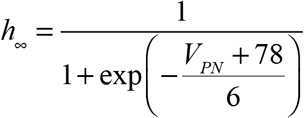. The time constants are given by 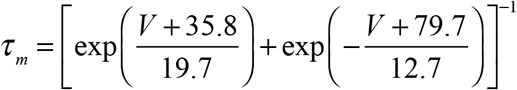 and 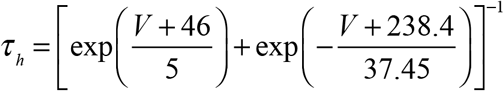. The inhibitory interneurons showed spike frequency adaptation due to a calcium dependent potassium current. The equations and parameter values are as follows,

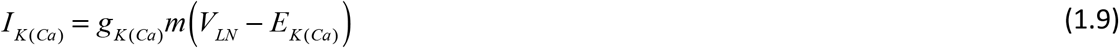

where, *g_K(Ca)_* = 0.3*μS*, and *E*_*K(Ca)*=−95*mV*_. The steady state value of the activation variable, *m*, is 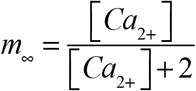 and the time constant is 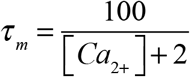. The calcium concentration, [*Ca*_2+_], dynamics is governed by the following equation,

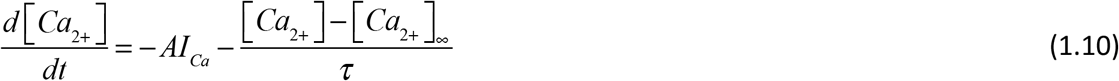

where the equilibrium concentration of Calcium, [*Ca*_2+_]_∞_ = 2.4 × 10^−4^*mM*, and the time constant *τ* = 5*ms*. The constant 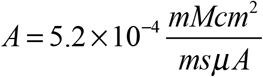. The calcium current in the inhibitory neurons is given by,

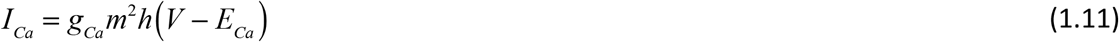

where, *g_Ca_* = 2*μS* and *E_Ca_* = 140*mV*. 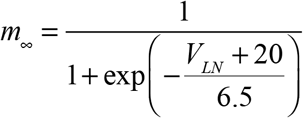 and 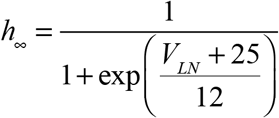. The time constants are *τ_m_* = 1.5 and 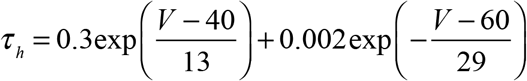

Fast GABAergic synapses between interneurons and between PNs and inhibitory interneurons were modelled using first order activation schemes. Similarly, nicotinic cholinergic input from PNs was used to drive the inhibitory interneurons. 50 of the 350 PNs extended excitatory input to the other PNs. All other PNs did not extend direct connections to each other. GABAergic and cholinergic synapses were both described by the following equations,

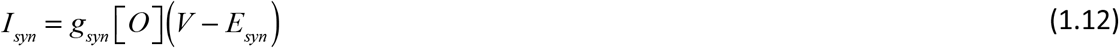

where the reversal potential is *E_nAch_* = 0*mV* for cholinergic receptors and *E_GABA_A__* = −70*mV* for fast GABA receptors. [*O*] is the fraction of open channels that is

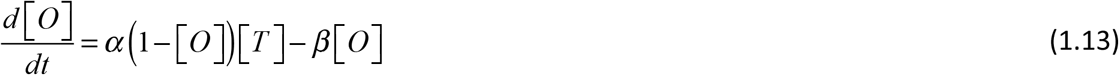

The rate constants, *α* = 10*ms*^−1^ and *β* = 0.16*ms*^−1^ for GABAergic synapses and *α* = 10*ms*^−1^ and *β* = 0.2*ms*^−1^ for cholinergic synapses. When the receptors are activated following a spike, the term [*T*] becomes non-zero. For cholinergic neurons this was modelled as the product of Heaviside functions in the following form,

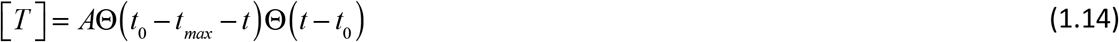

where, *t*_0_ is the time of receptor activation, *A* = 0.5 and *t_max_* = 0.3*ms*.

For GABAergic synapses,

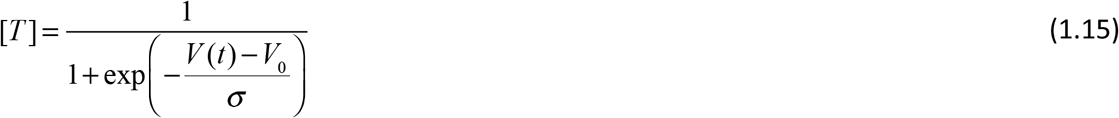

### Kenyon cells and MBONS

We modeled a large array (15000) of KCs and 100 MBONs. Given the large number of KCs, we modeled each as a two–dimensional map that can replicate in a computationally efficient way the dynamics of a variety of conductance–based neurons and networks of these neurons, but is computationally efficient (30,31). KCs and MBONs were modeled as regular spiking cells governed by the following equations,

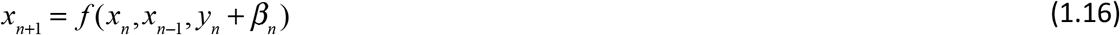

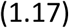

where the function *f* (*x_n_,x_n−1_,y_n_* + *β_n_*) is defined as,

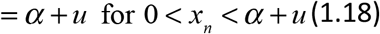

where *u* = *y_n_* + *β_n_ μ* = 0.0005 *σ* = 0.06 *β* = 0.03. Both KCs and MBONs received feedforward excitatory input. We did not model any lateral inhibition in these layers. Synaptic input is described by the following equations,

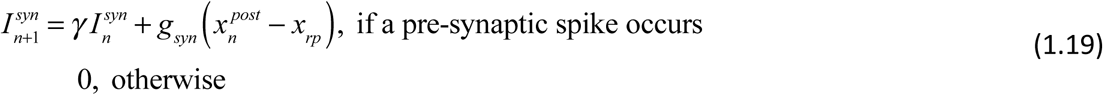

The KC-MBON synapse showed spike timing dependent plasticity. We modeled STDP using a model with an online update rule. Each pre-synaptic KC spike activated a variable that decayed exponentially post activation in the absence of other spikes. The dynamics of *x* followed the equation,

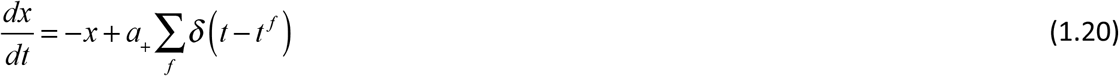

where, *t^f^* is the time at which a spike occurs. The effect of the spike on the weight of the KC-MBON synapse is given by the factor *a*_+_. In the absence of a spike the variable *x* decays exponentially to zero. A similar synaptic trace was defined to respond to postsynaptic spikes given by the following equation,

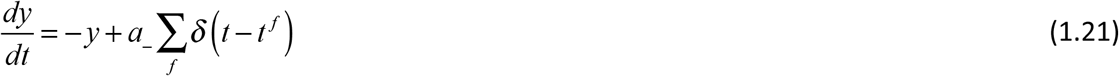

The weight of the synapse evolved in response to the timing of the pre- and post-synaptic spikes according to the following equation,

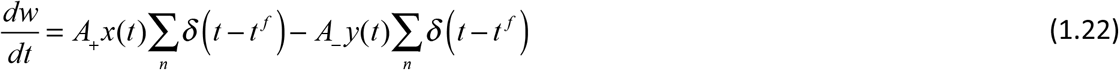

Further, the factors *A*_+_ and *A*_−_ changed in a manner that depended on the current weight *w*(*t*) of the synapse. Increases in weight when the synapse was close to a maximum weight were lower in magnitude than when it was further away from *W_max_*. This was achieved by introducing soft bounds to the weight by setting *A*_+_ = (*W_max_* − *w*)*η_+_* and *A*_−_ = *wη*_−_

## Acknowledgements

CA was funded by DBT–Wellcome India Alliance through an Intermediate fellowship IA/I/11/2500290 and IISER Pune. MB was supported by NIDCD grant (R01 DC012943)

